# Decreased RORC expression and downstream signaling in HTLV-1-associated Adult T-cell Lymphoma/Leukemia uncovers an antiproliferative IL17 link: a potential target for immunotherapy?

**DOI:** 10.1101/326900

**Authors:** Kritika Subramanian, Tim Dierckx, Ricardo Khouri, Soraya Maria Menezes, Lourdes Farre, Achilea Bittencourt, Keisuke Kataoka, Seishi Ogawa, Johan Van Weyenbergh

**Author notes:** Equal contribution (Kritika Subramanian, Tim Dierckx). **Corresponding Author: Johan Van Weyenbergh** Address: KU Leuven, Department of Microbiology and Immunology, Rega Institute for Medical Research, Clinical and Epidemiological Virology, B-3000 Leuven, Belgium.

## Abstract

Retinoic acid-related drugs have shown promising pre-clinical activity in Adult T-cell Leukemia/Lymphoma (ATL), but RORC (Retinoic acid Orphan Receptor C) signaling has not been explored. Therefore, we investigated transcriptome-wide interactions of the RORC pathway in Human T-cell Leukemia Virus-1 (HTLV-1) infection and ATL, using our own and publicly available gene expression data for ATL and other leukemias, HTLV-1-infected individuals and healthy controls. Gene expression data from ATL patients were analyzed using Weighted Gene Correlation Network Analysis (WGCNA) to determine gene modules and their correlation to clinical and molecular data. Both PBMCs and CD4+ T-cells showed decreased RORC expression in four different ATL cohorts. A small subset of RORChi ATL patients was identified with significantly lower pathognomonic CADM1 and HBZ levels but similar levels of other ATL markers (CD4/CD25/CCR4), hinting at a less aggressive ATL subtype. In addition, an age-dependent decrease in RORC expression was found in HTLV-1-infected individuals, but not in healthy controls, suggesting an early molecular event predisposing to leukemogenesis. Genes upstream of RORC signaling were members of a proliferative gene module (containing proliferation markers PCNA/MKI67), whereas downstream members clustered in an antiproliferative gene module. IL17C transcripts showed the strongest negative correlation to PCNA in both ATL cohorts, which was replicated in two large cohorts of T- and B-cell acute leukemias. In conclusion, decreased RORC expression and downstream signaling might represent an early event in ATL pathogenesis. An antiproliferative IL17C/PCNA link is shared between ATL, T-ALL and B-ALL, suggesting (immuno)therapeutic benefit of boosting RORC/IL17 signaling.

**Abbreviations:**

HTLV-1: Human T-cell Leukemia Virus-1
ATL: Adult T-cell Leukemia/Lymphoma
WGCNA: Weighted Gene Correlation Network Analysis
ATRA: All-trans retinoic acid
RARa: Retinoic acid receptor alpha
RORC: Retinoic acid orphan receptor C
TBLV: Type B Leukomogenic Virus
AML: Acute Myeloid Leukemia
T-ALL: T-cell acute lymphoblastic leukemia
IL: Interleukin
PCNA: Proliferating cell nuclear antigen
ID: Infectious Dermatitis

## Introduction

Human T-Lymphotropic Virus -1 (HTLV-1) is a retrovirus with an estimated prevalence of 10-20 million worldwide^1^. A recent return to the original name of Human T-cell Leukemia Virus-1^2^ is in agreement with its exceptional oncogenicity^3^. Although most HTLV-1 infections are asymptomatic, 2-6% of HTLV-1 infected individuals develop a CD4+CD25+ chemotherapy-resistant and aggressive leukemia known as Adult T-cell Lymphoma/Leukemia (ATL)^4–6^. ATL presents after a long latency period of the virus, commonly more than 20 years^7^. Patients therefore tend to be older individuals with an average age at diagnosis of 40 years in Central and South America and 60 years in Japan. Depending on the subtype (acute, lymphomatous, chronic, and smoldering), survival ranges from 4 months to over 5 years^8^.

HTLV-1 has two viral oncoproteins: Tax and HBZ. Tax benefits cell survival in HTLV-1 infected T-cells by interacting with NFκB^9^. However, Tax levels are undetectable in most ATL patients, either due to gene deletion or altered DNA methylation levels, whereas HBZ is expressed consistently in ATL^9^. HBZ modulates Tax expression and induces CD4+ T-cell proliferation^5^. CADM1/TSLC1 is also consistently expressed in ATL cells, such that CADM1 staining overlaps with the CD4+CD25+ T-cells in ATL and proviral sequences from these leukemic CD4+CADM1+ cells were consistently positive for the HBZ region^5^. Thus, CADM1 is a sensitive biomarker for ATL and might be used to determine treatment efficacy5/20/18 10:49:00 AM.

ATL patients display an increased incidence of opportunistic infections^10^, which could be attributed to a deregulation of the Th17 axis as an intact Th17 response is necessary for the clearance of opportunistic infections^11–14^. IL-17 and its upstream regulator IL-6 were increased in long-term cultured Tax+CD4+ T-cell supernatant at 24 and 72 hours^15^. IL-17 mRNA was also found to be highly expressed in HTLV-1 infected T-cells and Tax-expressing Jurkat cells^16^. Therefore, we hypothesize Tax-negative ATL cells are unlikely to express IL-17. Induction of the Th17 axis via retinoic acid receptors (RARs) and RAR-like orphan receptors (RORs) could potentially alleviate the increased opportunistic infection frequency caused by Th17 deregulation.

Retinoic acid blocks Th17 differentiation and stimulates regulatory T cell (Treg) production^17^. Although HTLV-1 proviral integration in the host genome showed greater enrichment of promotor sequence motifs binding p53 and STAT1 instead of the RORC locus^18^, downstream effects of p53 and STAT1 downregulate RORC transcription by suppressing STAT3^19^. The relevance of RORC in leukemogenesis is further supported by the observed increased proliferation and apoptosis rates in mice deficient in the protein product of the RORC gene ROR*γ*, leading to the development of T-cell lymphoma^20^ and lymphoblastic lymphoma^21^.

Taken together, deregulation of the RORC/Th17 axis can provide an explanation to both the oncogenic persistence of ATL and to patient susceptibility to opportunistic infections. In this study, we generate a representative consensus gene set for the RORC pathway of the Th17 axis and proceed to a systems biology analysis of novel and existing data to test the biological significance of this pathway in ATL.

## Methods

### In silico analysis

RORC expression levels were examined in publicly available transcriptomic data sets from patients with ATL, HTLV-1 infected asymptomatic controls, and healthy controls. A total of 135 untreated ATL patients, 12 HAM patients, 40 asymptomatic controls (AC), and 242 healthy controls (HC) from the Gene Expression Omnibus datasets GSE55851, GSE33615, GSE19080, GSE85487, and the European Genome-phenome archive EGAD1001411 dataset were used in this study (Table 1). EGAD1001411 initially contained 45 ATL patients, but one outlier with an overall strongly divergent transcriptome was removed. The effect of age on RORC expression was investigated in the Healthy Estonian Cohort for healthy controls (n=293) and the UK Cohort for HTLV-1 infected individuals (n=30).

**Table 1.** Transcriptomic (microarray and RNAseq) datasets used in RORC expression analyses

| Data set | Source | Cell Type | Disease Status | Sample Size |
| --- | --- | --- | --- | --- |
| GSE55851<br>Japanese Cohort #1 | Kobayashi et al. (2014) | CD4<sup>+</sup> T-cells | HC | 3 |
|  |  |  | AC | 6 |
|  |  |  | ATL | 12 |
| EGAD1001411<br>Japanese Cohort #2 | Kataoka et al. (2015) | CD4<sup>+</sup> T-cells | HC | 3 |
|  |  | PBMCs | AC | 3 |
|  |  |  | ATL | 44 |
| GSE33615<br>Japanese Cohort #3 | Yamagishi et al. (2012) | CD4<sup>+</sup> T-cells | HC | 21 |
|  |  | PBMCs | ATL | 52 |
| GSE19080<br>Caribbean Cohort | Oliere et al. (2010) | CD4<sup>+</sup> T-cells<br>(Immunoarray) | HC | 8 |
|  |  |  | AC | 11 |
|  |  |  | ATL | 7 |
|  |  |  | HAM | 12 |
| GSE85487<br>Brazilian Cohort | Dierckx et al. (2017) | PBMCs* | HC | 5 |
|  |  |  | ATL – Untreated | 8 |
| | | | ATL - IFN-α | 6 |
| | | | ATL - IFN-β | 6 |
| GSE29312<br>UK Cohort | Tattermusch et al. (2012) | Whole Blood | HC | 9 |
|  |  |  | AC | 20 |
|  |  |  | HAM | 10 |
| GSE78840<br>Healthy Estonian Cohort | Kasela et al. (2017) | CD4<sup>+</sup> T-cells | HC | 293 |
|  |  | CD8<sup>+</sup> T-cells | HC | 283 |
|  |  | PBMCs | HC | 77 |
| ImmuCo | Wang et al. (2015) | CD4<sup>+</sup> T-cells | HC | 551 |
|  |  | CD8<sup>+</sup> T-cells | HC | 149 |
|  |  | Bone marrow Mononuclear Cells | AML | 814 |
|  |  | Acute T-cell Leukemia | T-ALL | 138 |
|  |  | Acute B-cell Leukemia | B-ALL | 300 |

HC = Healthy Control. AC = Asymptomatic HTLV-1 Infected Control. ATL = Adult T-cell Lymphoma/Leukemia patients. HAM = HTLV-1 Associated Myopathy. AML = Acute Myeloid Leukemia. ALL = Acute Lymphoblastic Leukemia.

The Japanese Cohort #2 (EGAD1001411) RNA-Seq data was quality- and adapter-cleaned using trimmomatic^22^ and cutadapt^23^ and quantified with kallisto^24^ using an index built on the transcriptome obtained from the Genome Reference Consortium GRCh38, rel79. CIBERSORT was used to generate an in silico approximation of the relative composition of 22 immune cell types in the samples^25^.

To facilitate consistent analysis of both the microarray and RNA-Seq data, the ensemble and/or Agilent IDs of the datasets were matched with corresponding Entrez IDs using the biomaRt package^26,27^ in R. The Entrez IDs were verified with the associated GPL files on GEO where available. Considering transcriptomic analysis of the Caribbean Cohort was performed on a limited (non-genome-wide) microarray platform, 2134 Entrez IDs were common to all examined microarrays and comprised the list of genes examined in this study. To address the bias in the measurements inherent to each platform, we adapted the quantile discretization method proposed by Warnat et al.^28^ and transformed gene expression levels into percentile ranks among the surveyed 2134 genes for the meta-analysis. To further exclude the possibility of biasing our results, we refrain from making direct statistical comparisons of gene expression levels between datasets.

Published literature on RORc and ROR*γ*, as cited and detailed in the results section, was used to develop a consensus molecular pathway, which was then combined with the published ATL disease signature^29^ to develop a gene set for further exploration.

Weighted Gene Correlation Network Analysis (WGCNA)^30^ clusters genes into modules according to their topological overlap measure which quantifies how many gene-correlates were common to both members of each gene pair. To determine coherent gene modules and their correlation to clinical and molecular data, we performed WGCNA on each of the transcriptomic datasets from two independent ATL cohorts recently published by our group: *in vitro* gene expression data from short-term cultured ATL patient PBMCs (n=8, Brazilian Cohort) performed in parallel with lymphoproliferation, and *ex vivo* expression data from ATL patient PBMCs (n=44) of Japanese Cohort #2^29^. Module membership of the RORC gene set and the ATL signature genes were determined and correlated to demographic, clinical, and *in vitro* data.

### In vitro analysis

Spontaneous lymphoproliferation and apoptosis of primary cells (PBMC) from ATL patients (n=8, Brazilian Cohort) was measured by [^3^H]-thymidine incorporation, as described previously^31^.

### Statistical analysis

Statistical analysis was performed using GraphPad Prism 7.0. Differences in RORC gene expression were analyzed by Kruskall-Wallis test for Japanese Cohorts #1 and #2, and the Caribbean Cohort. For Japanese Cohort #3, where ACs were not included, Mann-Whitney was used to compare HC and ATL patients. The false discovery rate two-stage method of Benjamini, Krieger, and Yekutieli was used to correct for multiple comparisons. Spearman’s Rho was used to correlate gene expression (either per gene or per WGCNA module using their eigengene expression) to demographic (age), clinical data (patient survival) and *in vitro* data (proliferation and apoptosis).

## Results

### Transcriptomic analysis of four independent cohorts reveals a RORClo *ex vivo* phenotype in ATL

Gene expression profiling of *ex vivo* primary cells from ATL patients showed decreased RORC normalized expression in all four independent cohorts, revealing a common RORClo phenotype (Figure 1A-B-C-D). Japanese Cohort #3 (n=73) and Caribbean Cohort (n=38) had significant decreases in RORC expression of ATL patients (p<0.0001 and p=0.016 respectively). Japanese Cohorts #1 (n=18) and #2 (n=50) had borderline significant decreases in RORC expression of ATL patients (p=0.083 and p=0.10). HAM patients in the Caribbean Cohort did not have a significant change in RORC expression (p=0.54), however asymptomatic HTLV-1 infected individuals (AC) did display a significant decrease in RORC expression (p=0.016) when compared to healthy controls. ACs in other cohorts were not found to have a significant change in RORC expression, relative to healthy controls. Thus, RORC expression, measured as normalized expression (Fig. 1) and percentile rank (Suppl. Figure S1), is consistently lower in ATL than in HC, but varies among cohorts for AC. Since RORC gene expression had been previously shown to decrease in AC^32^, we performed a meta-analysis of the fold changes in RORC expression in all four cohorts, using both normalized expression and percentile ranks. Normalized gene expression allows for a comparison of the fold change in absolute RORC levels across cohorts but does not take into account the profound perturbation of the cellular transcriptome between healthy vs. leukemic CD4+ cells in AC and ATL patients, respectively. In contrast, percentile ranks are a measure of RORC expression relative to the overall transcriptome for each individual, which is more suitable for a comparison between divergent disease states. This was confirmed by two-way ANOVA, analyzing cohorts and disease status (HC-AC-ATL) as separate variables. For fold-change RORC normalized expression, cohort differences accounted for 21.2 % of variation (p=0.34) and disease status for 47.2 % of variation (p=0.06). For fold-change RORC percentile ranks, only 1.3 % of variation (p=0.48) was explained by cohorts and 95.8 % of variation (p<0.0001) was explained by disease status. As shown in Fig. 1E, RORC normalized expression was significantly (p<0.01) decreased in ATL patients, but not AC. Figure 1F displays the median RORC percentile rank fold-change, showing a significant 30% decrease in asymptomatic HTLV-1-infected individuals (p<0.0001, vs. HC) and an even further (43%) decrease in ATL patients (p<0.0001, vs. HC; p=0.006 vs. AC). This two-step decrease in RORC gene expression in ATL pathogenesis, first upon HTLV-1 infection and next upon progression to malignant disease, prompted us to investigate the possible influence of age upon RORC expression.

**Figure 1.**
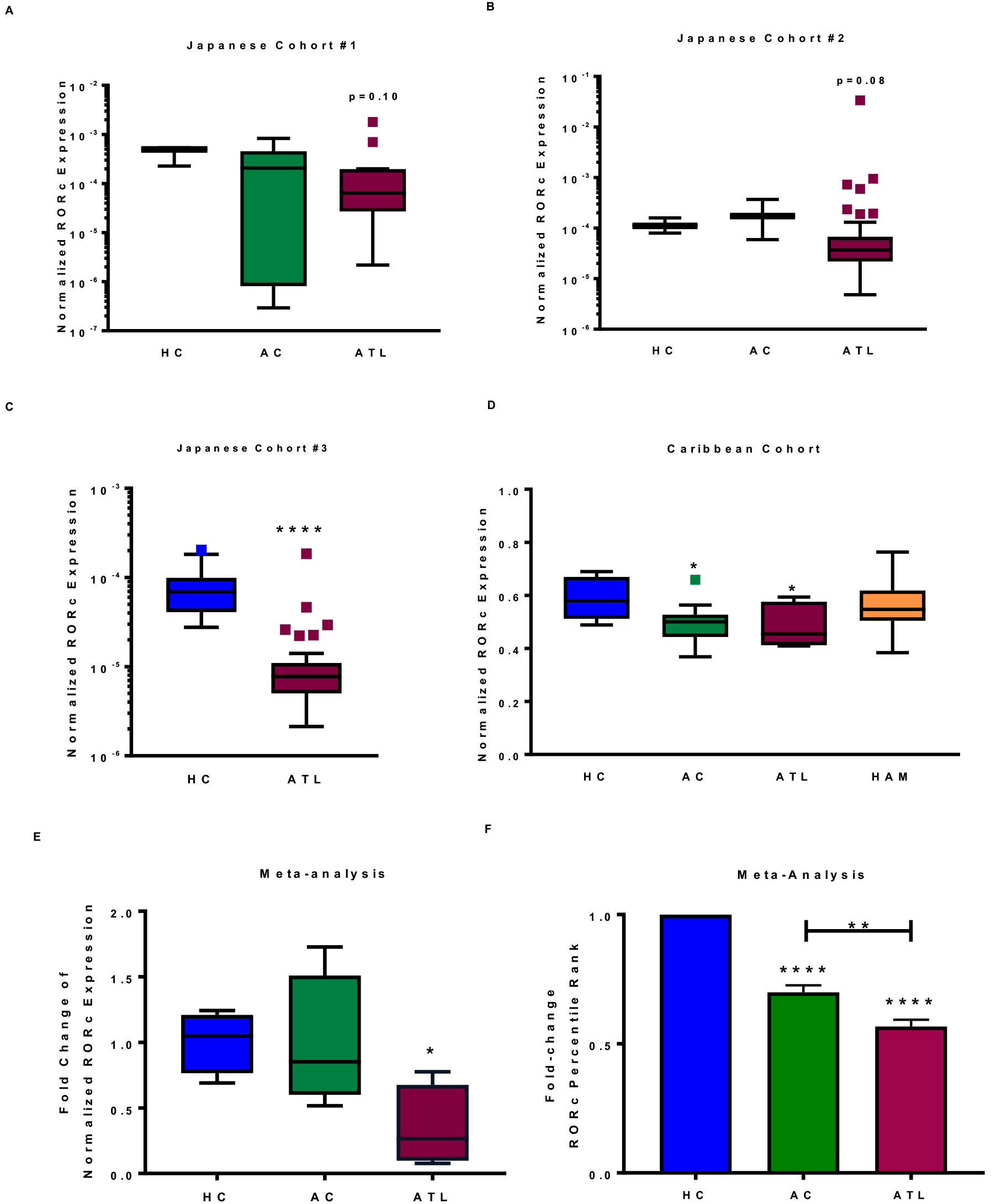
Normalized RORC expression levels for four independent cohorts consisting of ATL patients and healthy uninfected (HC) and/or HTLV-1 infected healthy controls (AC) showed a consistent decreased expression in ATL (A-D). (E) Meta-analysis of RORC normalized expression fold-change by disease status shows a significant decrease in ATL patients but not ACs. (F) Meta-analysis of RORC percentile rank fold-change by disease statusshows a significant two-step decrease for ACs and ATL. HCs = Healthy Controls, ACs = Asymptomatic Controls, ATL = Adult T-cell Lymphoma/Leukemia Patients, HAM = HTLV-1-Associated Myelopathy patients. *p<0.05, **p<0.01, ****p<0.0001.

### RORC Expression is not influenced by Age in Healthy Controls but decreases with Age in HTLV-1 Infected individuals

Since ATL usually occurs after several decades of HTLV-1 infection^5^ and ROR*γ*t Tregs were shown to increase with age in mice^33^, we investigated the effect of age upon RORC gene expression in healthy controls and HTLV-1-infected individuals from several cohorts. First, we examined paired CD4+ T-cells (n=293), CD8+ T-cells (n=283), and PBMC (n=77) microarray results from a cohort of healthy controls with sufficient power to study the effects of age (Healthy Estonian Cohort, Table 1). We found that RORC expression levels did not significantly change with age in CD4+ T-cells (r= 0.002, p= 0.45, Figure 3C), CD8+ T-cells (r=0.0001, p= 0.86, Figure 3D) or PBMC (r=0.0001, p=0.93), nor with gender (data not shown). In contrast, we found that RORC expression significantly decreased with age in HTLV-1 infected individuals without ATL, either AC and HAM/TSP patients (r=-0.57, p=0.0002, n=30 from UK Cohort, Figure 3A). We observed a similar tendency of decreased RORC expression with age (r=−0.62), but this observation did not reach statistical significance levels (p=0.10), most probably due to the small size of this ATL cohort (n=8, Brazilian Cohort) (Figure 3B). Unfortunately, the age of ATL patients was not available for the larger Japanese cohort.

### A minor RORChi subgroup of ATL patients displays a unique CADM1loHBZlo phenotype

RORChi outliers (Rout Method^34^, Q=0.1%) were observed in the three Japanese cohorts, accounting for a total of 13 out of 108 ATL patients (12.04%), Therefore, we examined this phenotype more closely in the largest examined cohort (Japanese Cohort #2), where 7 outliers with a higher normalized RORC expression were identified (Figure 1B). The patients from this cohort were then split into two groups, according to their RORC levels, as shown in Figure 2. Interestingly, we noted that RORC expression was inversely associated with expression levels of pathognomonic ATL markers CADM1 and HBZ. Thus, RORChi patients displayed significantly lower HBZ (p=0.0061) and CADM1 (p=0.045) levels, but similar expression levels of other ATL surface marker genes (*CD4*, *CD25/1L2RA*, *CCR4*) suggesting the RORChi subgroup might represent a distinct, possibly clinically relevant, molecular subgroup of ATL. The lower CADM1 and HBZ expression levels in RORChi patients may represent the decreased proliferation rate of chronic or less aggressive ATL subtypes. As shown in Supplementary Figure S2, RORChi patients showed similar expression levels of other ATL driver genes (*STAT3*, *PLCG1*, *NFKB1*, *RELA*, *FAS*)^29,35^, highlighting the specificity of the RORChiCADM1loHBZlo phenotype. Positive expression of IRF4 and c-REL has been associated with resistance to IFN-α + AZT therapy in ATL patients^35,36^. Interestingly, IRF4 and c-REL expression did not differ between RORChi and RORClo patients (Suppl. Figure S2). This finding suggests RORC expression is independent of IFN-α + AZT therapeutic resistance and offers an additional molecular target for patients failing this therapy.

**Figure 2.**
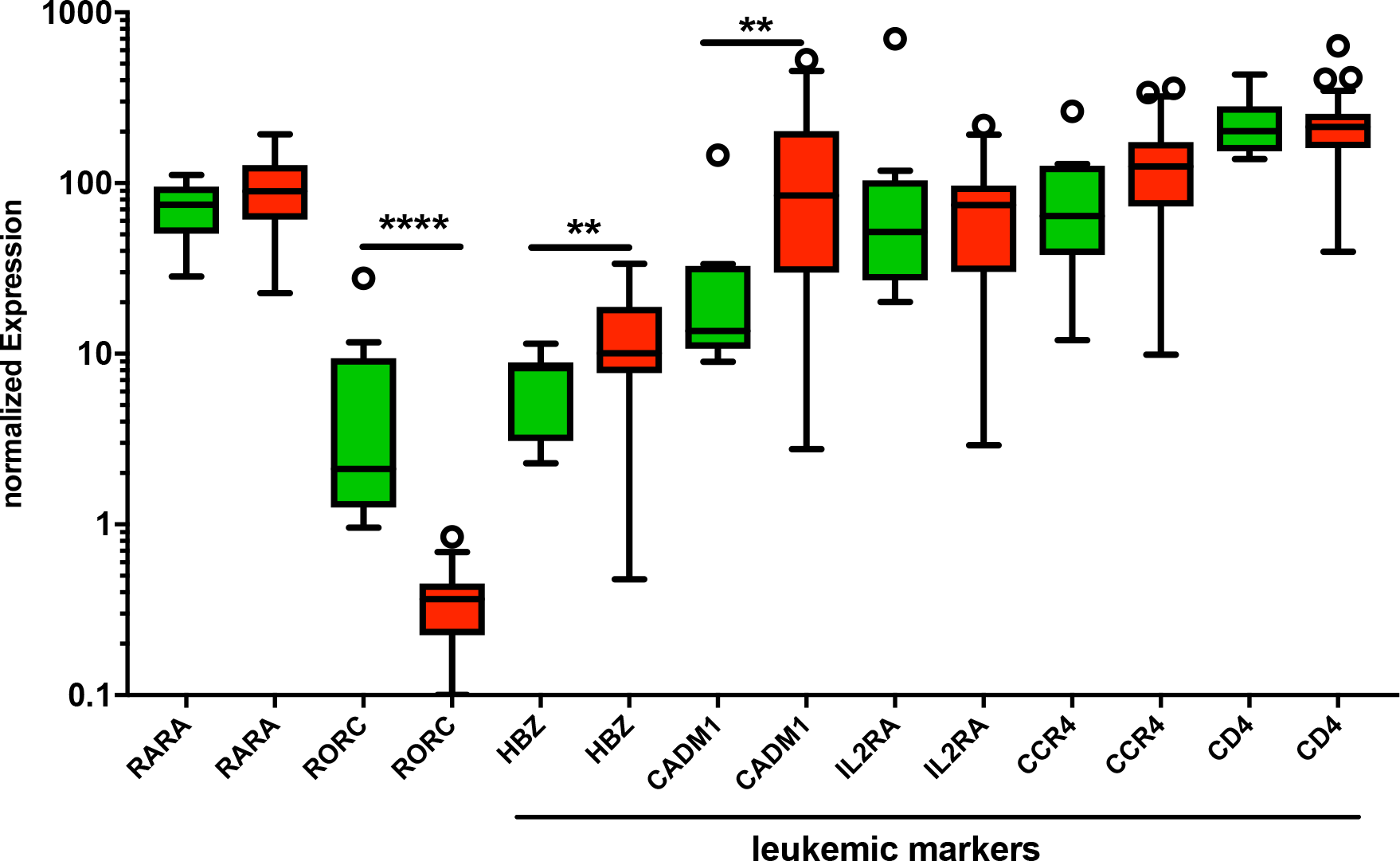
RORC expression levels of ATL patients from Japanese Cohort #2 separated into two groups: RORChi (Green) and RORClo (Red) show RORChi levels were associated with lower HBZ and CADM1 expression levels. RORChi (7 outliers) and RORClo groups were compared with expression levels for ATL driver/mutated genes. *p<0.05 **p<0.01 ****p<0.0001

**Figure 3.**
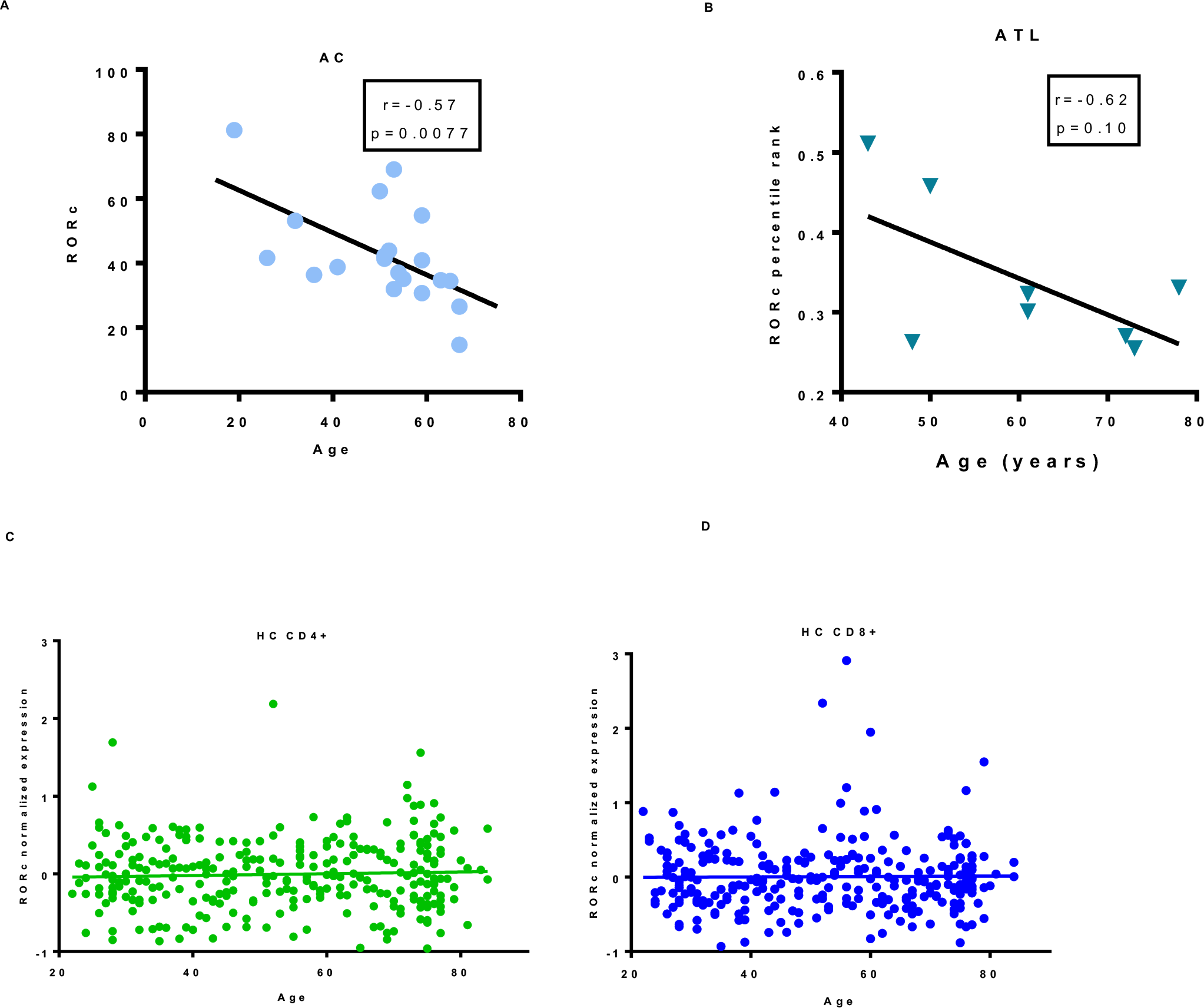
HTLV-1 infected Individuals from UK Cohort (A) and ATL patients from Brazilian Cohort (B) showed a decrease in RORC expression as age increased. This decrease was absent in healthy controls, in either CD4+ cells (C), CD8+ cells-(D) or PBMCs (not shown).

### Definition of a consensus RORC pathway and gene set and its relevance to ATL oncogenesis

To facilitate the molecular exploration of the RORChi phenotype, a RORC gene set was determined based on published literature findings, integrating the intrinsic oncogenic pathway for STAT3 activation, as defined by Yu et al.^37^, and RARA/RORC signaling summarized by Muranski and Restifo (2013)^38^. The consensus RORC pathway included IL6, IL23, IL21, IRF4, BATF, STAT1, STAT5, RARa, TGFβ, NF**κ**B, SLC2A1 (GLUT1), BCL6, STAT3, FOXP3, SOCS1, RORC, and IL17A/F. Figure 4 illustrates the interplay between these genes, as detailed in the legend. RNX1, T-bet, RORA, and TGFB1R were not measured by the microarray used for the initial WGCNA analysis on the Brazilian cohort (pilot cohort) and therefore excluded from the gene set.

**Figure 4.**
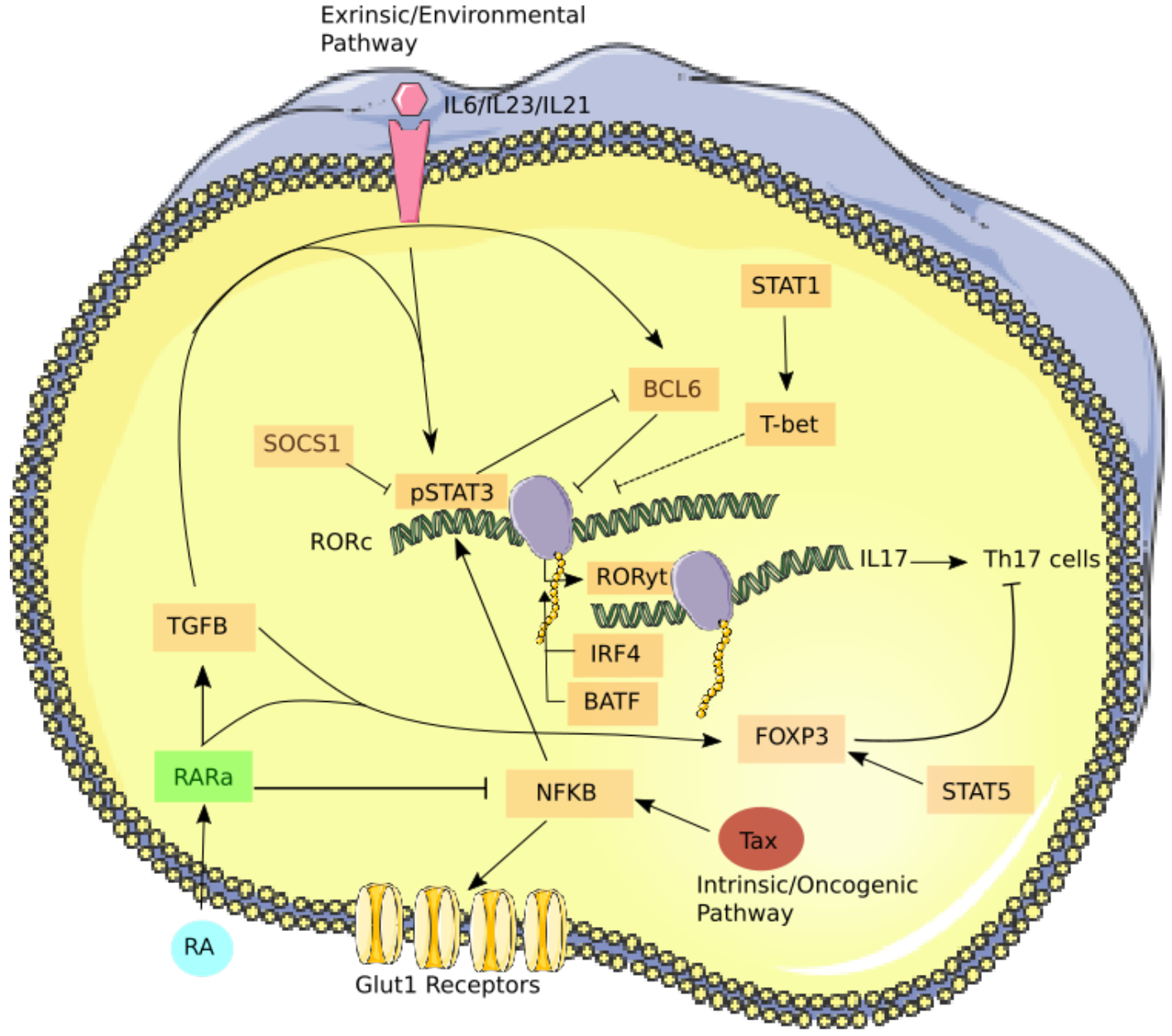
Simplified figure depicting the roles of RORC Pathway members, adapted from Yu et al. (2009)^36^ and Muranski and Restifo (2013)^37^. The figure was produced using Servier Medical Art (http://www.servier.com) and edited using Inkscape software.

Transcriptomic expression levels of RORC pathway members extracted from a UK HTLV-1 infected asymptomatic control dataset (UK Cohort; GSE29312) and ATL cohort (Japanese Cohort #2; EGAD1001411) showed that the majority were expressed at highly variable levels (Supplementary Figure S3). First, prominent STAT1 expression is in line with published findings in AC^32,38^ and ATL^35,39,40^. On the other hand, downstream members of the RORC pathway and particularly, IL17 family genes were either undetectable or poorly expressed.

WGCNA analysis of PBMCs from our pilot ATL cohort (n=8, Brazil, Figure 5A) showed overlap of the RORC pathway with a gene module correlated to proliferation, containing bona fide proliferation markers *PCNA* (Proliferating Cell Nuclear Antigen) and *MKI67* (the gene coding for Ki67 antigen, routinely used in flow cytometric quantification of proliferation). As shown in Figure 5B, downstream pathway members RORC and the IL17 family, underexpressed in ATL, were positively correlated with the proliferative module and negatively correlated with the antiproliferative module. Similarly, upstream and overexpressed gene members of the RORC pathway displayed the reverse trend. This resulted in a significant bifurcation in the RORC pathway when correlation coefficients of member genes from the antiproliferative module were regressed with the correlation coefficients from the proliferative module (Supplementary Figure S4, r=0.97, p<0.0001). Overall, WGCNA analysis suggested that inducing RORC and suppressing upstream pathway members may decrease the cell proliferation rate in ATL.

**Figure 5.**
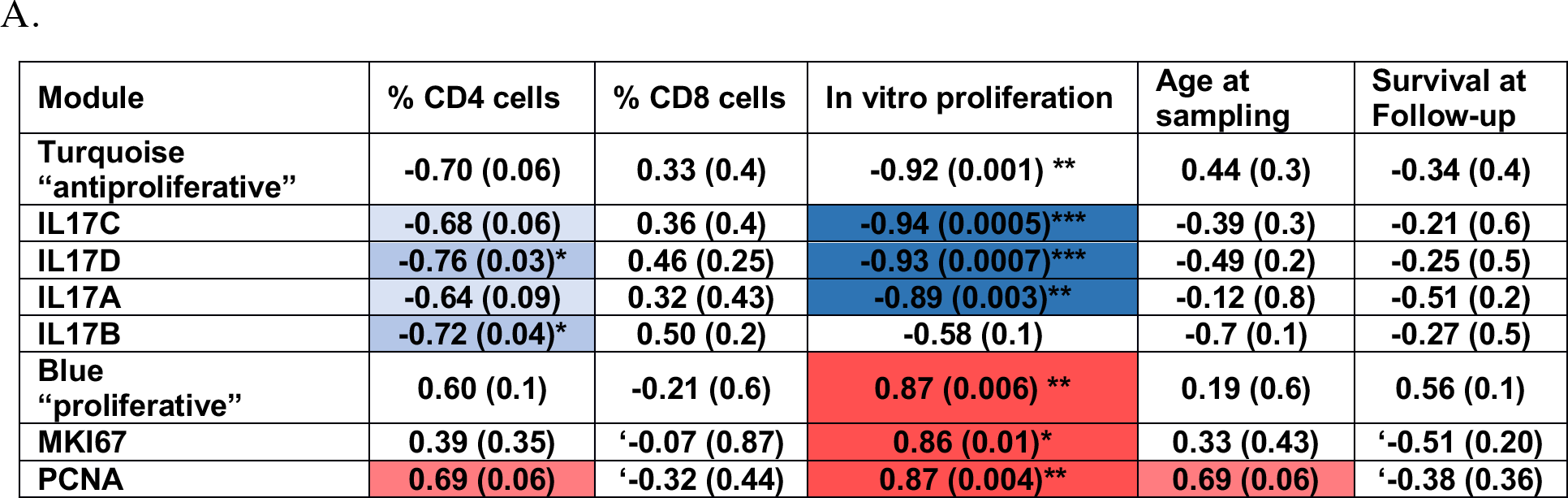
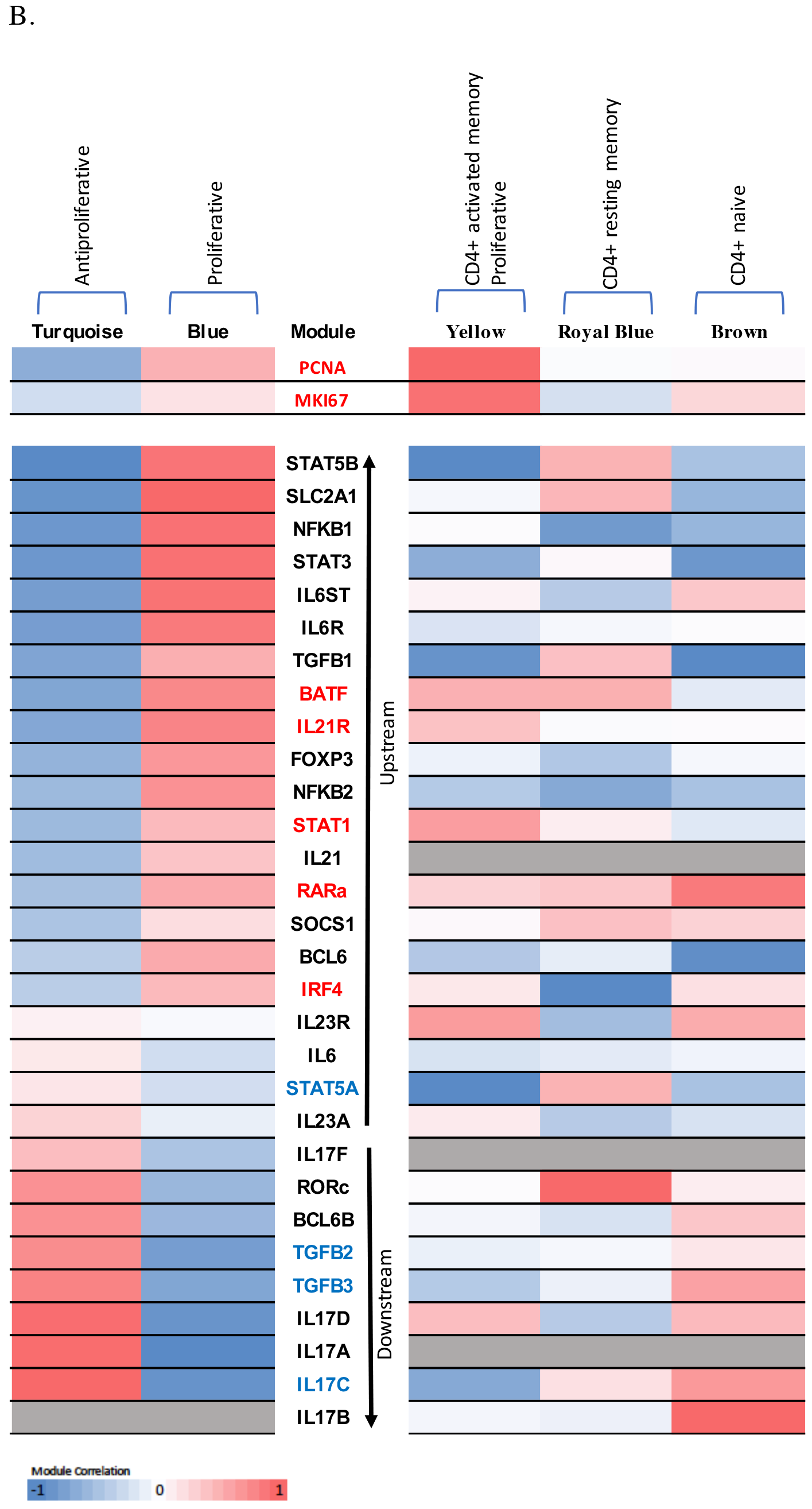
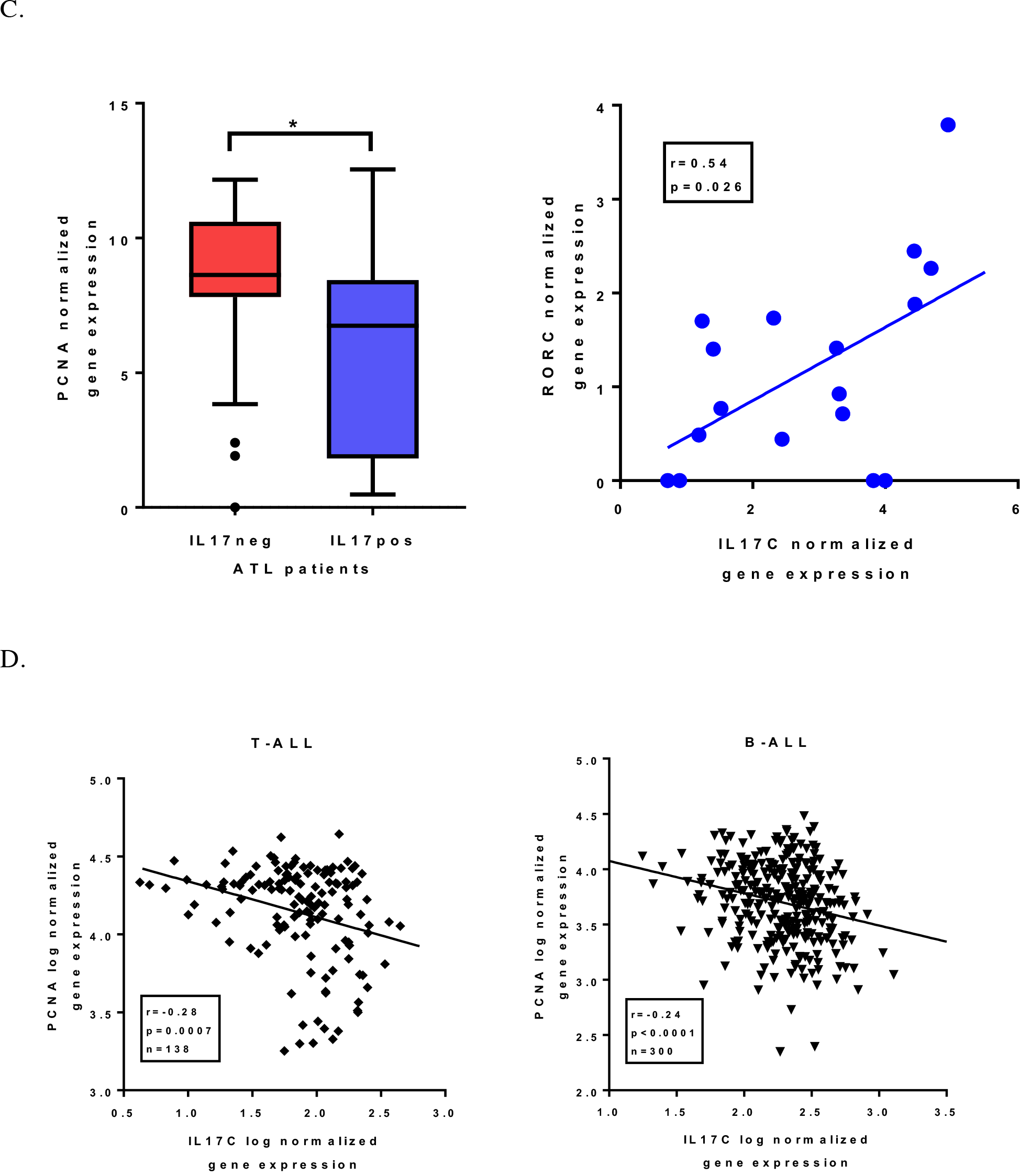
Modular transcriptomic analysis of primary ATL cells reveals a strong association of the RORC/IL17 pathway with proliferation. A) WGCNA findings for the Turquoise and Blue modules are shown for selected molecular and clinical correlations. R-value and p-values (in parenthesis) are shown *p<0.05 **p<0.01. B) Most RORC pathway members were positively correlated with the turquoise (“antiproliferative”) gene module in primary ATL cells and negatively correlated with the blue (“proliferative”) modules, such that most downstream members were found to be associated with the antiproliferative module (Brazilian cohort, n=8, left panel). WGCNA of Japanese Cohort #2 (right panel, n=44) shows RORC has a positive correlation with CD4+ resting memory T-cells (Royal Blue module, r=0.47, p=0.001). Yellow module is positively correlated with CD4+ memory active T-cells (r=0.39, p=0.007), as well proliferation markers PCNA (r=0.92, p=10^−5^) and MKI67 (r=0.87, p=10^−2^). Brown module is positively correlated with CD4+ naïve T-cells (r=0.49, p=0.0006). Genes which were validated in both ATL cohorts for proliferative modules are colored according to their R-values. C) ATL patients expressing IL17C (IL17pos, n=17, Japanese Cohort #2) showed decreased PCNA expression as compared to patients with undetectable IL17C (IL17neg, n=28), and IL17C levels were positively correlated with RORC levels. D) A negative correlation between IL17C and PCNA transcript levels was replicated in large T-ALL (n=138) and B-ALL cohorts (n=300).

To confirm and extend these findings on proliferation, we repeated the WGCNA in the larger cohort of ATL patients (n=44, Japanese cohort #2). We additionally obtained *in silico* estimates of the relative size of 22 immune cell type populations using the CIBERSORT software^25^. As shown in Figure 5B, RORC was the only pathway member which was significantly and positively correlated (r=0.42) with the presence of resting memory CD4+ T-cells (p=0.0041). Downstream pathway members IL17B (r=0.62, p= 0.0000054) and IL17C (r=0.42, p=0.04) were positively correlated with the presence of naïve CD4+ T-cells. STAT3 inducer NFκB subunits 1 and 2 were negatively correlated with naïve CD4+ T-cells (p=0.02 and p=0.052 respectively) and resting memory CD4+ T-cells (p-0.000073 and p=0.00084 respectively). Similar to the observations in the WGCNA of the pilot cohort, a reverse trend was also seen in the CIBERSORT analysis between upstream and downstream members of the RORC pathway and their correlation with naïve and activated memory CD4+ T-cell fractions (Figure 5B). Together, the two WGCNA analyses, combined with CIBERSORT CD4+ subtype quantification, suggest a distinct change in proliferative pathways between upstream and downstream members of the RORC/IL17 pathway with opposite effects in activated memory vs. naïve and resting memory CD4+ T-cells. Among downstream pathway members, IL17C showed the strongest antiproliferative gene module membership in both cohorts and was also more frequently detected than other IL17 family members (IL17A/B/D/F). Therefore, we classified ATL patients from the largest cohort (Japanese cohort #2) into IL17C expressing, (IL17Cpos, n=17) and IL17C negative (IL17Cneg, n=28). As shown in Figure 5C, IL17C positive patients had significantly lower gene expression levels of proliferative marker PCNA (Mann-Whitney p=0.022) and in those patients, IL17C was positively correlated to RORC gene expression (r=0.54, p=0.026), confirming the findings of our modular analysis. To explore if the antiproliferative IL17C/PCNA link might be specific to ATL or shared with other leukemias, we analyzed two large cohorts of acute T- and B-cell leukemia (T-ALL, n=138; B-ALL, n=300). Similar to ATL, we found a significant negative correlation between IL17C and PCNA transcript levels in both T-ALL (r=−0.24, p=0.007) and B-ALL (r=−0.28, p<0.0001).

### RORC expression is independent of viral HBZ and Tax mRNA

SLC2A1, STAT3, and NFkB1 are members of the proliferative module (Figure 5) and upstream to RORC in the RORC pathway (Figure 4), of which NF**κ**B1 and STAT3 also belong to the mutational signature of ATL^29^. Therefore, we tested WGCNA module membership of two major HTLV-1 transcripts: HBZ and Tax, which are, respectively increased and decreased in ATL cells compared to asymptomatic controls^39–42^. However, no overlap was found between the module memberships of RORC, HBZ, and Tax in the WGCNA of Japanese Cohort #2 (data not shown). In addition, no RORC gene module members were significantly correlated to HBZ or Tax transcript levels, suggesting decreased RORC levels and signaling in ATL are not a direct consequence of retroviral transcription.

### IFN-α, IFN-β and Ascorbic Acid *in vitro* treatment differentially regulates RORC expression in primary ATL cells and HTLV-1 transformed cell lines

We previously tested the effects of IFN-α and Ascorbic Acid on HTLV-1 infected transformed cell lines (MT2, MT4, C8166)^41–43^. Although both these drugs have shown efficacy in decreasing HTLV-1-induced proliferation^41–44^, only the high-dose AA affected the retinoic acid pathway, specifically the shared RORC/Th17 pathway. Reanalysis of our transcriptomic data showed that neither IFN-α nor high-dose Ascorbic Acid altered RORC expression levels (log fold-change = 0.042, p= 0.59 and log fold-change = 0.069, p= 0.39, respectively). Ascorbic acid stimulated an increase in expression of a key gene in Th17 differentiation, IL23R (log fold change = 0.81, p=0.000024), in support of its possible use in (combination) therapy for ATL. RARa expression was unchanged by IFN-α (log fold change = 0.003, p=0.98) but decreased by ascorbic acid (log fold change = −0.28, p= 0.064). Interestingly, our *in vitro* data (Brazilian cohort) demonstrated that RARa levels are upregulated upon *in vitro* treatment of ATL PBMCs with IFN-β (log fold-change = 0.31, p=0.017), but not IFN-α. IFN-β was more likely to significantly alter the expression levels of STAT1, IRF4, TGFB1R, IL23R, FOXP3, and IL6, while IFN-α significantly altered BCL6 only, as shown in Supplementary Figure S5. This is in agreement with our recently demonstrated differential anti-proliferative and pro-apoptotic effect of both IFN subtypes^31^.

## Discussion

Upon transcriptomic meta-analysis of four different cohorts, we found a specific and consistent RORClo phenotype in primary ATL cells and to a lesser extent in HTLV-1-infected individuals, in contrast to healthy controls. In addition, HTLV-1-infected individuals displayed an age-dependent decrease in RORC expression. The observed two-step decrease of RORC in ACs and ATL patients might thus represent an early event in HTLV-1-driven leukemogenesis. We also identified a small subset (12.04%) of RORChi ATL patients with significantly lower pathognomonic CADM1 and HBZ levels but similar levels of other ATL markers (CD4, CD25 and CCR4), hinting at a less aggressive ATL subtype.

ATL pathogenesis develops over decades as is seen by patients presenting at least 20 years after HTLV-1 infection; yet not all infected patients develop ATL. Observational studies suggest that ATL, at least in the Caribbean and Brazil, can be triggered by the pediatric cutaneous manifestation known as Infectious Dermatitis^45–48^. ID is a chronic, eczematous condition with scaly, crusted lesions often superimposed by *Staphylococcus aureus* or *Streptococcus pyogenes* infections^45,46^. Defects in the Th17 axis increase vulnerability to *S. aureus* and *Candida albicans* infections, whereas *in vivo S. aureus* primed memory Th17 cells inhibited IL-17 production and increased IL-10 production^47,49^. Of interest, two recent papers have demonstrated a role for IL-10 as an unexpected proliferative trigger of infected CD4+ T-cell clones and, possibly, leukemogenesis^50,51^. Corroborating these findings, IL10 was found to be a significant (r=0.36, p=0. 013) member of the proliferative module, together with PCNA and MKI67 in our WGCNA analysis. Therefore, our findings underscore an IL-10 vs. RORC/IL-17 antagonism in HTLV-1-associated pathologies provides a possible molecular basis for the epidemiological link between ID and ATL, alteration of the RORC/Th17 axis, and subsequent progression to leukemogenesis. Indeed, modular transcriptomic analysis in ATL shows a strong correlation of the RORC pathway with cell proliferation and possibly oncogenesis, which underscores its therapeutic potential. WGCNA analysis combined with CIBERSORT suggested the involvement of RORC pathway members in the homeostasis of resting memory and naïve CD4+ T-cells. Combining the RORClo observation in ATL cohorts with our WGCNA analysis, we find that decreased RORC expression is correlated with proliferation and ATL driver genes (STAT3, NF-kB). Thus, inducing RORC and switching to a RORChi phenotype may convert ATL cells to a less aggressive subtype, suggested by the lower CADM1 and HBZ levels seen in the RORChi subset (Figure 2). Inducing IL17 expression via RORC stimulation would also subsequently alter the host immune response to reduce the risk of opportunistic infections by increasing Th17 cell count. In addition, the strongest negative correlation observed in both ATL cohorts, between IL17C and proliferation marker PCNA, was replicated in two large cohorts of other acute lymphoid leukemias, namely TALL and B-ALL (Figure 5D). This indicates an antagonistic regulation between Th17 cells, usually considered as pro-inflammatory, and leukemic cell proliferation. Regarding the clinical translation of these results, antitumor immunotherapy using Th17 cells has recently shown promising results in animal models. Adoptive cell therapy using *ex vivo* Th17 cell selection enhanced antitumor activity^50,52^, to a greater extent than Th1 cells and other CD4+ T-cells^50,52^.

Although most often believed to antagonize IL17 production, IFN-β can trigger and even exacerbate IL17 production, especially in Th17-mediated inflammatory diseases^51,53^. This becomes problematic in cases of multiple sclerosis, where 30-50% of patients are resistant to IFN-β therapy^51^. However, this same exacerbation could be useful in ATL as a means of increasing Th17 cell production and decreasing proliferation of leukemic clones. IFN-β significantly alters the expression of more RORC pathway members than IFN-α (Supplementary Figure S5), a common therapeutic adjuvant to zidovudine in ATL treatment. This finding, along with the observation that IFN-β has superior anti-proliferative and pro-apoptotic properties compared to IFN-α^31^, makes IFN-β a novel, valuable option for combination therapy in ATL.

Recently, immune checkpoint inhibitors have come to the forefront of anticancer immunotherapy^52,53^. Studies have suggested that inhibiting the transmembrane protein Programmed death ligand −1 (PD-L1) can increase Th17 cell count. Notably, human anti-PD-L1 antibodies restored IL-17A protein levels in naïve T-cells of patients with a loss-of-function STAT3 mutation^52–54^. Also noteworthy is the induction of Th17 cell differentiation by RORy agonist LY C-54143, which simultaneously reduced PD-1+ cell numbers and PD-1 expression *in vitro*, resulting in tumor growth inhibition in two murine models^55^. PD-L1 amplifications have been associated with worse prognosis in ATL patients, especially in aggressive subtypes^56^.

In conclusion, we describe a predominant RORc^lo^ phenotype observed in four cohorts of ATL patients and a minor RORChi molecular subgroup with significantly lower pathognomonic CADM1 and HBZ mRNA levels. An age-dependent decline in RORC level indicates a possible early event in HTLV-1-driven leukemogenesis, supported by modular transcriptomic analysis of ATL patients, revealing a strong negative correlation of the RORC/IL17 pathway with proliferation, which was shared with T-ALL and B-ALL patients. Thus, inducing RORC levels and/or signaling might represent (immuno)therapeutic benefit in ATL and possibly other acute lymphoid leukemias.

## Supplementary Figure Legends

**Supplementary Figure S1.** RORC percentile ranks for four independent cohorts consisting of ATL patients and healthy uninfected (HC) and/or HTLV-1 infected healthy controls (AC) showed a consistent decrease in expression in ATL (A-D). HCs = Healthy Controls, ACs = Asymptomatic Controls, ATL = Adult T-cell Lymphoma/Leukemia Patients, HAM = HTLV-1-Associated Myelopathy patients. *p<0.05, **p<0.01, ***p<0.001, ****p<0.0001.

**Supplementary Figure S2.** Differential RORC expression of ATL patients from Japanese Cohort #2 is independent of ATL driver genes and genes involved in IFN/AZT response.

**Supplementary Figure S3.** RORC and downstream IL17 family genes are poorly expressed in both asymptomatic HTLV-1 infected individuals (top) and ATL patients (bottom).

**Supplementary Figure S4.** RORC and downstream IL17 family genes cluster in an antiproliferative gene module. Using WGCNA analysis, a significant bifurcation in the RORC pathway is observed when correlation coefficients of member genes from the antiproliferative module (Turquoise) were regressed with the correlation coefficients from the proliferative (Blue) module (r=−0.97, p<0.0001) in primary cells from ATL patients (Brazilian cohort).

**Supplementary Figure S5:** Differential effects of IFN-β vs. IFN-α upon RORC pathway members in primary ATL cells. Blue represents genes which had significantly (p<0.05) altered expression levels with IFN-β treatment. BCL6 (Red) was the only gene member which was significantly altered by IFN-α treatment.

